# A cognitive representation in primary visual cortex modulated by vision

**DOI:** 10.64898/2026.08.29.748029

**Authors:** Kyu Hyun Lee, Philip Adenekan, Fan Gao, Jenny Zhou, Jose Hernandez, Allison Yorita, Razi Haque, Kenneth Kay, Massimo Scanziani, Loren M. Frank

## Abstract

Primary visual cortex (V1) is a critical substrate for mammalian vision. Traditionally, visual inputs are thought to be the main drivers of V1 activity, with internal signals playing a modulatory role. Here we show that this relationship is inverted for a large fraction of V1 neurons. In rats completing a navigation task in darkness, these neurons encoded progress along physically distinct paths with a shared turn structure. Under illumination, visual stimuli gain-modulated this path-invariant activity rather than replacing it with stimulus-driven responses.

Path-invariant V1 neurons were also preferentially coordinated with hippocampal ensembles during sharp-wave ripples, linking them to a brain-wide network involved in learning. These findings establish that an internal model of the world can serve as a primary driver of activity in sensory cortex.

## Main Text

Sensory information guides behavior, but does not determine it. Adaptive behavior requires combining sensory input with internal representations of cognitive variables, such as where the animal is and what it is trying to achieve. A visual landmark, for example, is useful to a navigating animal not simply because of its visual features, but because those features can update an estimate of progress through a learned path. How sensory information is integrated with such internal representations remains poorly understood.

Primary visual cortex (V1) provides a natural setting in which to address this question. As the first cortical stage of visual processing, V1 is classically associated with retinotopic maps and low-level visual features such as orientation and direction of motion (*1*, *2*). Yet studies have shown that nonvisual variables can drive V1 activity, even in the absence of visual input. These include other sensory modalities, such as auditory (*3*), somatosensory (*4*), and vestibular (*5*) signals, as well as movement-related variables, such as locomotion (*6*, *7*) and movements of the head (*8–10*), eye (*11*), and face (*12*, *13*). Beyond these sensory and motor signals, V1 is also shaped by cognitive variables, including attention (*14*), task context (*15–17*), predictions about visual input (*18*, *19*), and spatial position (*20–22*). Much of this work on cognitive variables in V1 has focused on how they modulate visually evoked activity (*14*, *20*, *21*, *23*, *24*). Yet whether V1 can explicitly represent a cognitive variable in the absence of visual input, and how such a representation interacts with vision, is unknown.

Freely moving navigation in a structured environment provides a way to address these questions. Navigation engages rich cognitive representations of space and task structure, as exemplified by work in hippocampal-entorhinal circuits (*25*, *26*). Animals can perform learned navigation tasks in complete darkness, eliminating visual input while preserving task structure. The maze geometry can be chosen to differentiate between spatial and other abstract representations, such as progress along a path. Restoring or rearranging visual landmarks can then reveal how visual input transforms the ongoing internal representation, while simultaneous recordings from hippocampal area CA1 can test whether this representation is coordinated with a brain region known to encode spatial and task structure (*20*, *27*).

## A structured navigation task with controlled visual input

We therefore recorded large populations of neurons simultaneously from V1 and hippocampal area CA1 as rats navigated a W-shaped, three-armed maze (“W-maze”) in controlled visual conditions (Fig. 1A). We implanted four 128-channel high-density flexible polyimide neural probes (fig. S1A), with one probe targeting each hemisphere of V1 and CA1. Across four animals, we recorded >1,200 V1 neurons (mean ± SD: 316.5 ± 156.4 simultaneously recorded) and >400 CA1 neurons (mean ± SD: 107.3 ± 50.5 simultaneously recorded). Probe locations were verified histologically after the experiments (fig. S1A).

**Fig. 1.**
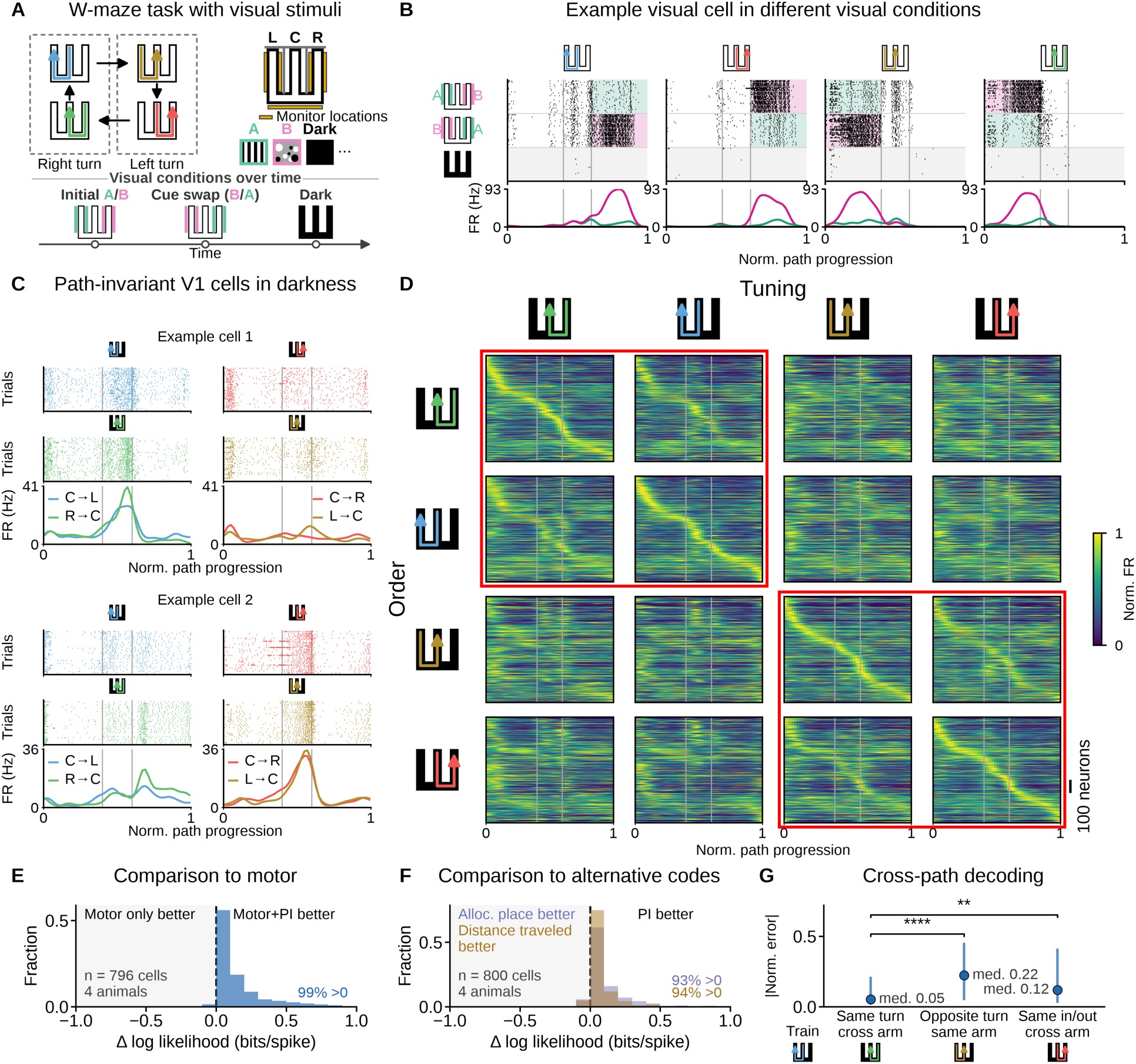
Path-invariant firing in V1 in darkness. (**A**) Freely moving rats performed a spatial alternation task on a W-maze with visual stimuli. Left: Four path types (two right-turn paths and two left-turn paths) that the animal alternates between to receive reward at the arm ends. Right: visual stimuli and their locations on the maze. Monitors cover areas shown as yellow bars (center arm is not covered). Bottom: Experimental progression. (**B**) A visually responsive V1 cell in three visual conditions (AB, BA, and darkness). Rasters and occupancy-normalized tuning curves are plotted against normalized path progression; gray lines mark path-segment boundaries and shading marks stimulus locations. The cell prefers stimulus B and is nearly silent in darkness. (**C**) Two V1 cells tuned to path-invariant progression, with similar responses on physically distinct same-turn paths in darkness. (**D**) Population tuning in darkness. For each row, neurons were ordered by peak position on the indicated path using half the trials; held-out tuning is shown across all four paths. Off-diagonal sequences within the red boxes indicate generalization across same-turn paths. (**E**) Held-out comparison of motor-only and motor plus path-invariant progression (PI) models; positive ΔLL favors motor+PI. (**F**) Held-out comparison of PI with allocentric-place (blue) and distance-traveled (red) models; positive ΔLL favors path-invariance. (**G**) A decoder trained on one path was tested on same-turn/cross-arm, opposite-turn/same-arm, or same- inbound-outbound/cross-arm paths. Points and bars show median absolute normalized error and interquartile range; lower values indicate better decoding. Trial-based permutation tests: **** *p*<10^-4^; ** *p*<10^-2^.

Rats were trained to perform a spatial alternation task on the W-maze. Starting from the center arm, animals were rewarded for visiting the outer arms in alternation and returning to the center arm. Correct performance thus consisted of four path types: center-to-left, left-to-center, center-to-right, and right-to-center (Fig. 1A). Each day of recording included at least two light epochs and a single dark epoch. In a light epoch, visual stimuli (black and white dots or sinusoidal gratings) were displayed on monitors that flanked the left and right arms as well as the horizontal segment of the maze. The presented stimuli were then swapped across the two outer arms in a subsequent light epoch to dissociate stimulus identity from location on the maze. In the dark epoch, all light sources were removed from the environment except for diffuse near-IR (940 nm) illumination used to track the animal’s behavior. The irradiance of the near-IR source was below published thresholds for visual detection in rats ((*28*, *29*); Methods). All analyses in this study used data from trained animals with performance > 90%.

## V1 encodes a cognitive variable in darkness

As expected, V1 activity differed strongly across the visual conditions. Approximately 27% of V1 neurons behaved like classic visual feature detectors: they were tuned to locations with specific visual stimuli (Fig. 1B) and were minimally active in darkness (<0.5 Hz, fig. S1B). In contrast, the remaining 73% of V1 neurons were active in darkness at this threshold (fig. S1B), and many showed reliable firing patterns across repeated traversals of the four paths on the maze (fig. S1C; fraction with split-half tuning correlation >0.5: V1 0.6, CA1 0.7). Although previous V1 recordings in virtual reality have revealed location-dependent modulation (*20–22*), the firing we observed was not tied to a single physical location. Instead, it was often similar across physically distinct paths that shared an overall turn direction (i.e., left vs. right). For example, a neuron active near the middle of a center-to-left path was also active near the analogous location of a right-to-center path (Fig. 1C). We refer to this selective firing at specific locations within paths and turn-specific invariance across paths as *path-invariant progression tuning*.

To visualize the population-level structure of this representation, we computed and plotted the path progression tuning curves of V1 neurons, ordered by their preferred location in each path. Neurons tiled the full path progression axis on each path, producing diagonal sequences in the tuning-curve matrix, and this ordering generalized across paths with the same turn direction, producing off-diagonal sequences (Fig. 1D). Thus, V1 population activity in darkness signals path-invariant progression.

Previous work has reported modulation of V1 by kinematic (e.g., speed, head movement (*7*, *9*, *10*)) and task-related (e.g., distance traveled (*24*)) variables. We therefore asked whether these variables could account for the activity observed in darkness. Because similar movements occurred at different physical locations and distances traveled, the task allowed us to compare these explanations with path-invariant progression.

A model that included path-invariant progression contributed predictive information beyond measured motor behavior. We fit nested Poisson GLMs and evaluated predictive performance on held-out data. The motor model included six variables: speed, acceleration, head direction, signed angular velocity, angular speed, and head angular acceleration. An expanded model included the same motor variables together with path-invariant progression. The expanded model improved held-out log likelihood for 99% of dark-active V1 neurons (median ΔLL = 0.08 bits/spike; Fig. 1E; per-animal data in fig. S2A). Thus, path-invariant progression contributed predictive structure broadly across the V1 population beyond that captured by the measured motor variables.

The path-invariant progression model was also better than models that relied on allocentric place or distance traveled. The allocentric place model captured tuning to physical location on the maze without directionality. The distance-traveled model captured linearized distance from the starting reward port across all four path types. The path-invariant progression model captured normalized progress through a path with shared tuning across paths with the same turn direction. On held-out data, the path-invariant model predicted activity better than the allocentric place model (93% positive ΔLL, median ΔLL = 0.05 bits/spike) and the distance-traveled model (94% positive ΔLL, median ΔLL = 0.03 bits/spike; Fig. 1F; per-animal data in fig. S2B). Thus, among the candidate task variables tested, path-invariant progression provided the most predictive description of V1 tuning.

Finally, we directly quantified path-invariance using cross-path generalization decoding. A Bayesian decoder trained on one path predicted the animal’s progress on a physically distinct same-turn path more accurately than on paths with the opposite turn direction or the same inbound/outbound type relative to the center arm (median absolute error normalized by path length: 0.05 vs. 0.22 vs. 0.12; Fig. 1G; per-animal data in fig. S2C). This transfer across physically distinct same-turn paths shows that V1 encodes path-invariant progression in darkness.

## Visual stimuli gain-modulate path-invariant progression tuning

We next asked how visual input affected the path-invariant progression tuning, first at the level of single neurons. Introducing visual stimuli often preserved the location of V1 firing within individual paths, while enhancing or suppressing its amplitude (Fig. 2A). For tuning curves spanning multiple path segments, these amplitude changes could differ across segments, consistent with the different visual stimuli encountered in each. They could also differ between the two same-turn paths that the V1 neurons exhibited invariance in darkness. Again, this was consistent with the fact that the animal traversing the two paths now experiences two different sequences of visual input.

**Fig. 2.**
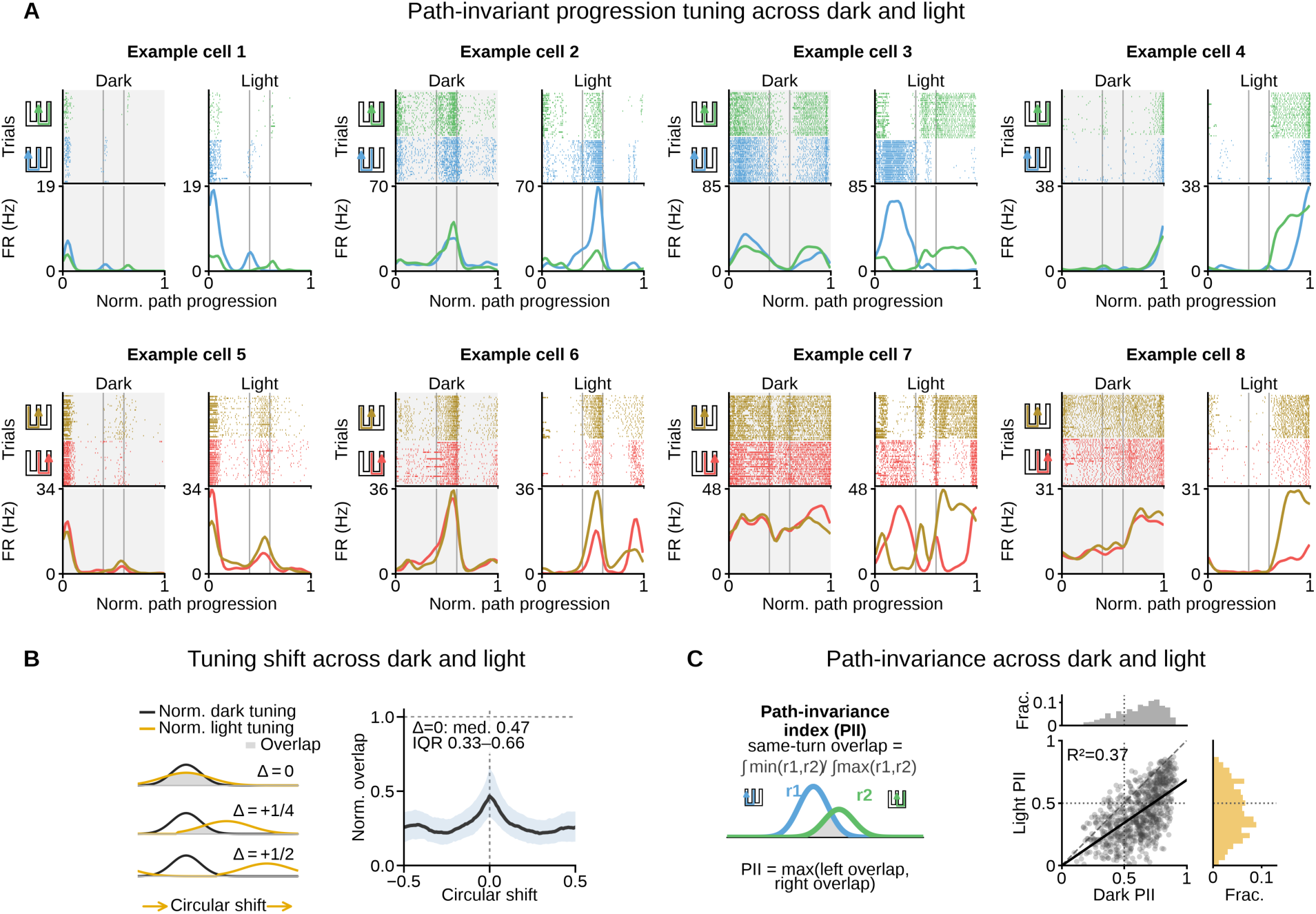
Visual input modulates path-invariant progression tuning in V1. (**A**) Raster plots and tuning curves for eight V1 neurons on same-turn path pairs in darkness and light. Cells 1–4 show the right-turn pair; cells 5–8, the left-turn pair. Colors denote paths; gray lines mark path-segment boundaries. (**B**) The light tuning curve was circularly shifted relative to the dark curve, and normalized overlap was measured at each shift. Population overlap peaked at zero shift (median, line; interquartile range, shading), indicating no systematic shift in tuning location. (**C**) Path-invariance index (PII) was defined as the larger of the tuning-curve overlaps for the left- and right-turn path pairs. Light PII was evaluated for the same turn direction selected in darkness. Dark and light PII values are correlated, but PII was lower in light. Black line, linear fit; dashed line, unity.

We quantified the stability of tuning by measuring the preservation of tuning location independently of peak rate. Specifically, we calculated the normalized overlap between each neuron’s dark and light tuning curves within individual paths while circularly shifting the light curve relative to the dark curve. The overlap peaked at zero shift, confirming that neurons tended to fire at the same locations in darkness and light (Fig. 2B; per-animal data in fig. S3A). After subtracting the overlap expected under circular misalignment and normalizing by the noise- ceiling, the dark-light overlap at zero shift reached a median of 0.47 of the explainable range across neurons (IQR, 0.33–0.66). Thus, a substantial degree of within-path progression tuning was preserved across lighting conditions.

We further quantified path-invariance using a gain-sensitive, unnormalized overlap between the tuning curves on physically distinct paths sharing turn direction. We refer to this as the path- invariance index. For each neuron, we calculated the path-invariance index for the same turn-pair in darkness and light.

Dark and light path-invariance indices were positively correlated (Fig. 2C; per-animal data in fig. S3B), indicating that neurons with strong path-invariance in darkness also tended to show strong path-invariance in light. Nevertheless, path-invariance indices were lower in light for many neurons, indicating that the tuning on same-turn paths became less similar when visual stimuli were present.

This reduction in path-invariance was also evident at the population level. Decoders trained on one path transferred less accurately to its same-turn partner in light than in darkness, whereas path-specific place decoding improved in light (fig. S4A; per-animal data in fig. S4C). Effective signal dimensionality was also higher in light than in darkness (fig. S4B), consistent with visual input introducing path-specific degrees of freedom into the lower-dimensional, path-invariant activity expressed in darkness. These differences were unlikely to arise from gross changes in behavior, as measured motor variables were similar across lighting conditions (fig. S5A-B).

Thus, visual input preserved substantial progression tuning within individual paths while differentiating the representations of paths that shared a common structure in darkness.

One mechanism that could jointly explain these effects is stimulus-specific gain (i.e., multiplicative) modulation of the dark tuning curves. Scaling the dark tuning by different gains in segments containing different stimuli would preserve its tuned location within each path while reducing similarity across same-turn paths and increasing path specificity at the population level.

Under a fixed visual configuration, however, stimulus identity covaried with location along each path, so this could not distinguish multiplicative gain from alternative explanations in which local visual responses replaced or added to the dark tuning. We therefore directly compared these three hypotheses on a held-out light epoch in which visual stimuli were swapped across the outer arms (AB to BA; Fig. 3A).

**Fig. 3.**
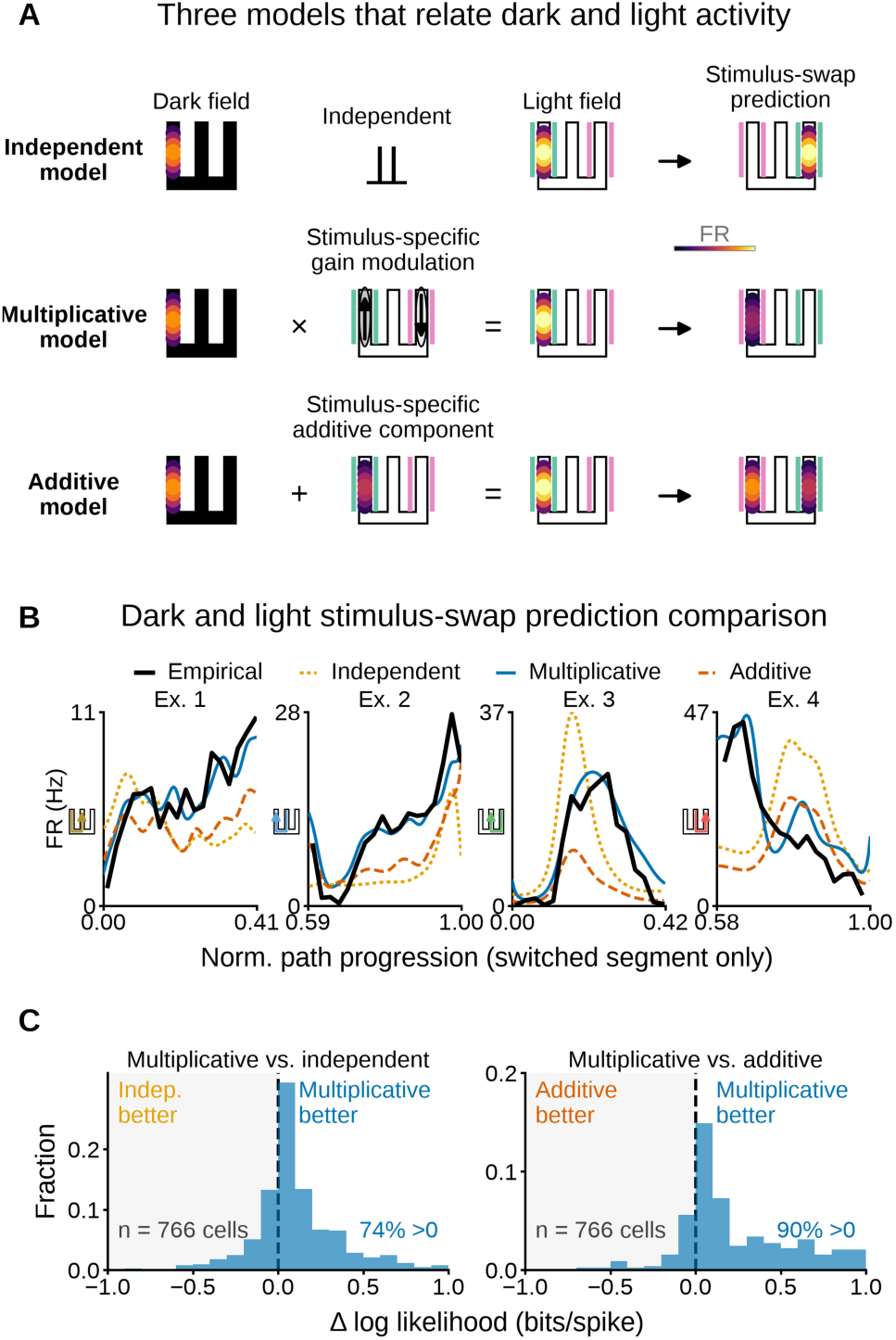
Multiplicative gain modulation best explains the effect of visual input on path- invariant progression tuning. (**A**) Three models relating dark and light activity. The independent model fits separate dark and light fields. The multiplicative model generates the light field by applying stimulus-specific gain (up and down arrows) to the dark field. The additive model generates the light field by adding a stimulus-specific component to the dark field. The rightmost column shows each model’s prediction for the stimulus-swap condition. **(B)** Empirical tuning curves (black) and predictions from the independent (yellow), multiplicative (blue), and additive (orange) models on the swapped segment of four example neurons. **(C)** Held-out log-likelihood differences in the stimulus-swapped light epoch for multiplicative vs. independent (left) and multiplicative vs. additive (right) models, averaged across the four paths. Positive ΔLL favors the multiplicative model.

To do so, we implemented the three candidate models and derived their divergent predictions on the held-out light epoch (Fig. 3A). The independent and multiplicative models were implemented as Poisson GLMs fitted to spiking during one dark epoch and one light epoch (Fig. 3A). The independent model estimated separate tuning curves in darkness and light, allowing the light response to follow the location of visual stimuli independently of the dark response. The multiplicative model instead constrained dark and light activity to share a single tuning curve and represented light-specific changes as a single visual-gain term for each of the three segments that made up a path. This model predicted that V1 activity would remain in the same location but be scaled up or down in the held-out light epoch. Finally, the prediction from the additive model was generated by estimating the light and dark tuning curves, taking their difference in one arm, and adding this to the dark tuning curve in the swapped arm.

We found that the multiplicative model was a better fit for most neurons. Despite using the fewest parameters, it predicted held-out light activity better than the independent or additive model for most dark-active V1 neurons (vs. independent: 74% positive ΔLL; median ΔLL = 0.07 bits/spike; vs. additive: 90% positive ΔLL; median ΔLL = 0.90 bits/spike; Fig. 3B–C). This advantage was consistent across animals and sessions (fig. S6A–B). In a three-way comparison, the multiplicative model outperformed both the independent and additive models in most neurons (54% preferring multiplicative vs. 27% independent and 19% additive; fig. S6C). Thus, for the largest fraction of neurons, visual input scaled path-invariant progression tuning from darkness rather than replacing it or adding a separate visual response.

## CA1 ripple ensembles are coordinated with path-invariant V1 neurons

The path-invariant activity we observed in V1 most likely arises from brain regions previously reported to express similar representations, such as hippocampal–entorhinal (*25*, *26*), prefrontal (*30*, *31*), and posterior parietal circuits (*32*, *33*). Activity across these regions is known to be coordinated during hippocampal sharp-wave ripple (SWR) events, brief population bursts seen during “off-line” periods (sleep and awake immobility) that contribute to memory storage and updating (*34–36*).

We therefore asked whether path-invariant neurons were preferentially coupled to hippocampal activity during SWRs. We first examined mean ripple-triggered firing rates in V1 and CA1. As expected, CA1 neurons showed strong modulation around hippocampal ripples (fig. S7A-B; per-animal data in fig. S7D). In contrast, mean firing-rate modulation in V1 was weak (fig. S7A-B). Thus, V1 engagement during ripples was not apparent as a uniform population response.

However, each SWR recruits a specific set of active CA1 neurons, and averaging activity across SWR events could obscure ensemble-specific coordination. The representational content of SWRs is known to be coordinated across hippocampus and cortical areas (*37*, *38*), raising the possibility that V1 coupling depends on which CA1 neurons are active in each SWR event.

Consistently, SWR-restricted cross-correlograms revealed that individual CA1 neurons were co-active with distinct subsets of V1 neurons (Fig. 4A). We further quantified this relationship using cross-validated Poisson GLMs that predicted each V1 neuron’s spike count in the 0–200 ms following ripple onset from the simultaneous CA1 population spike-count vector (Fig. 4B-C).

**Fig. 4.**
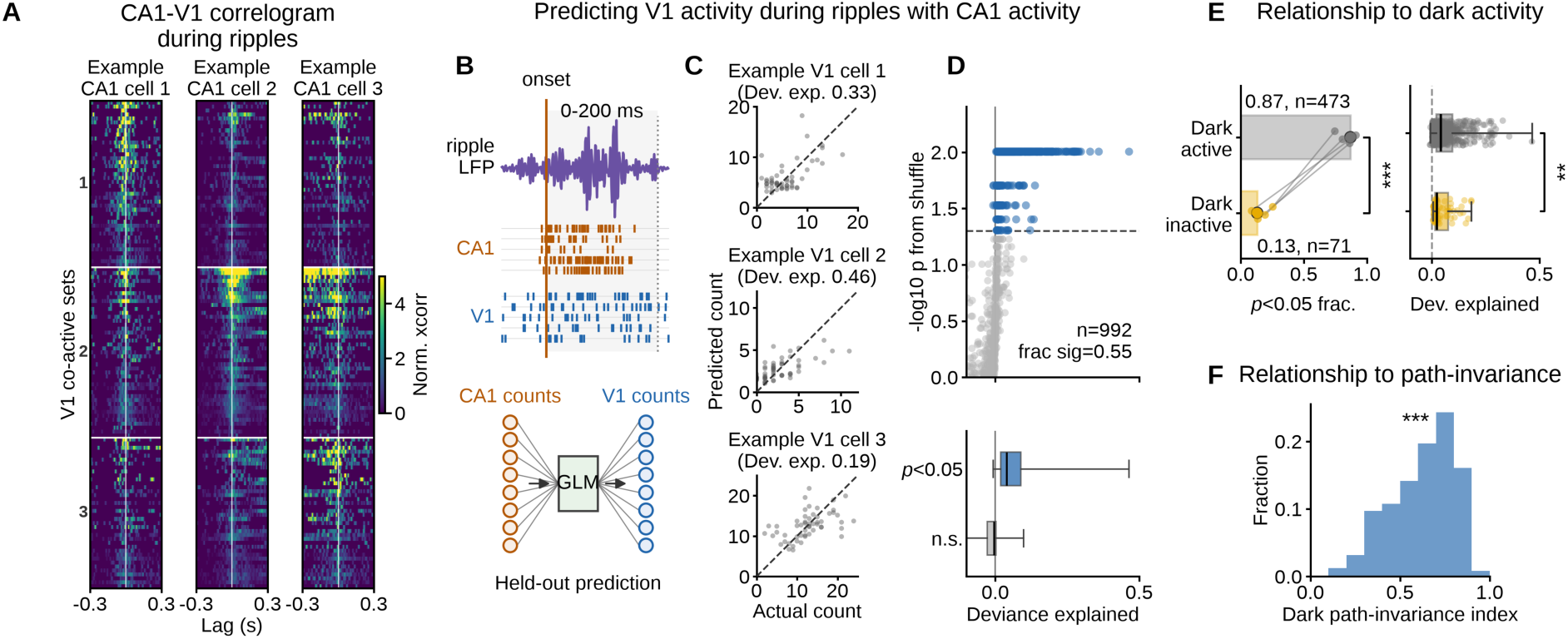
CA1 activity predicts path-invariant V1 neurons during hippocampal ripples. (**A**) Ripple-triggered cross-correlograms for three example CA1 neurons and simultaneously recorded V1 neurons. For each CA1 neuron, V1 neurons are grouped into three co-active subsets. (**B**) A Poisson GLM was used to predict each V1 neuron’s spike count in the 0–200 ms window after ripple onset from simultaneously recorded CA1 spikes (5-fold CV). **(C)** Held-out predicted versus actual ripple spike counts for three example V1 neurons. **(D)** Prediction performance for all V1 neurons with appreciable firing during SWRs (>0.1 spikes/ripple). Top, deviance explained and shuffle-test significance; dashed line, *p*=0.05. Bottom, deviance explained for significant (*p*<0.05) and nonsignificant groups. **(E)** SWR-predictable V1 neurons were more likely to be dark-active. Left, fraction of dark-active and dark-inactive neurons among SWR- predictable V1 cells. Right, deviance explained was higher for dark-active than dark-inactive neurons. (**F**) Dark-active, SWR-predictable V1 neurons had elevated dark path-invariance index (defined in Fig. 2C) relative to dark-active, non-SWR-predictable V1 neurons. For (E) and (F), significance was assessed by shuffling the relevant binary labels within each animal, preserving per-animal group sizes, and refitting the linear mixed effects model for each of 100,000 permutations (two-sided tests); \*\**p*<10^-2^, \*\*\**p*<10^-3^.

We found that CA1 activity significantly predicted firing in 55% of all V1 neurons with appreciable activity (>0.1 spikes/ripple) during held-out SWRs, relative to a null generated by permuting CA1–V1 pairings across events (Fig. 4D; per-animal data in fig. S8A). This permutation disrupts the correspondence between CA1 and V1 activity, and thus the results reflect ripple-by-ripple covariance. For these predictable V1 neurons, the full CA1 population vector also outperformed mean CA1 activity alone, indicating that the identity of the active CA1 ensemble carries predictive information over the global ripple strength (fig. S7C; per-animal data in fig. S7D).

SWR predictability was also preferentially associated with the dark-active, path-invariant V1 neurons. The SWR-predictable population was strongly enriched for dark-active neurons (87% vs. 13% dark-inactive; Fig. 4E; per-animal data in fig. S8B). Among predictable neurons, predictive performance was higher for dark-active than dark-inactive cells (Fig. 4E; per-animal data in fig. S8B). Furthermore, the dark-active neurons that were SWR-predictable showed higher path-invariance index than ones that were not SWR-predictable (Fig. 4F; per-animal data in fig. S8C). Thus, the V1 neurons most strongly coordinated with CA1 ripple ensembles included a substantial population carrying path-invariance in darkness. These results indicate that path-invariant V1 neurons were tightly coupled to specific CA1 neurons during SWRs, demonstrating engagement in a network critical for memory storage related to abstract representations.

## Discussion

In summary, we show that a cognitive variable related to path-invariant progression dominates V1 activity during free navigation in darkness. Visual inputs often gain-modulate this cognitive representation, inverting the standard account of V1 activity. V1 neurons with strong path-invariant tuning were also preferentially coupled to distinct CA1 ensembles during SWRs, demonstrating coherent network activity across V1 and CA1 consistent with top-down modulation of these neurons.

Previous studies have shown that cognitive variables shape V1 activity. Of particular relevance is work showing modulation of V1 activity by the animal’s location in virtual reality. For example, identical stimuli evoked different visual responses at different positions, and this modulation tracked experience, engagement, subjective position, and spatially specific expectations (*20–22*). These studies establish that visual responses in V1 are modulated by navigation-related internal variables.

In contrast, we probed the visual cortex of freely behaving animals in real-world environments with and without visual stimuli. Surprisingly, this experimental design revealed that a cognitive variable can be explicitly represented independently of visual input. Visual stimuli multiplicatively modulated this tuning in a stimulus-specific manner while preserving the underlying structure expressed in darkness. This suggests that the path-invariant progression tuning may provide a task-state scaffold on which visual inputs act, allowing visual stimuli to distinguish otherwise equivalent states on different paths. A similar organization may allow sensory evidence to act on cognitive variables in other sensory cortices.

Because path-invariance is defined by the learned environmental layout and rules of the task, its expression in V1 is unlikely to be innate. It must either be learned within V1 or supplied by other circuits, possibly maintained through interactions across a broader cognitive network. The strong coordination of path-invariant V1 neurons with CA1 during SWR events may thus reflect both the underlying anatomical pathways that enable communication across these circuits and a mechanism for learning and maintaining path-invariant representations in V1.

## Supplementary figures

**fig. S1.**
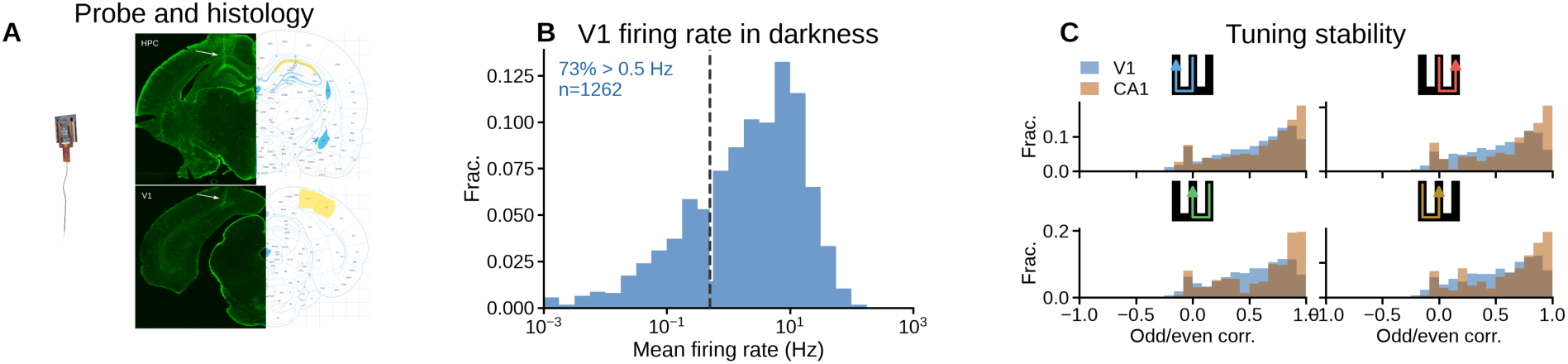
**(A)** Left: Flexible polyimide probe used in this study. Right: histology showing the implantation locations in dorsal hippocampus CA1 (top) and V1 (bottom). White arrows in GFAP staining (green) mark the probe tracks. **(B)** Distribution of mean firing rate of V1 neurons in darkness during the W-maze spatial alternation task. Only movement periods (>4 cm/s) are included. Dashed line marks 0.5 Hz, used as threshold for defining dark-active vs -inactive groups. **(C)** Distribution of stability of spatial tuning (defined as the correlation between tuning curves learned from even and odd trials) in darkness in each of the four path types, for both V1 (blue) and CA1 (orange).

**fig. S2.**
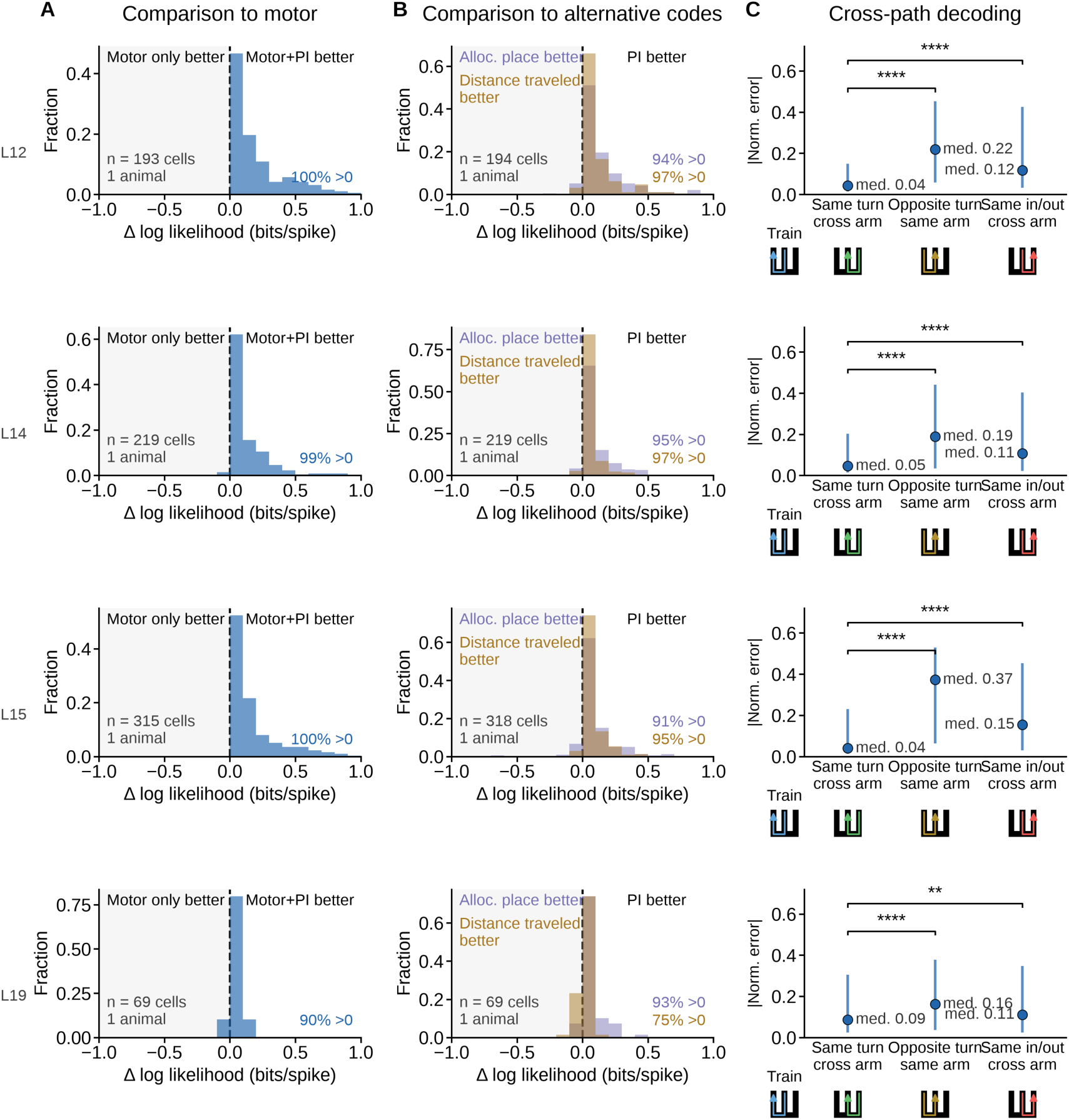
**(A)** Per-animal data of Fig. 1E. **(B)** Per-animal data of Fig. 1F. **(C)** Per-animal data of Fig. 1G.

**fig. S3.**
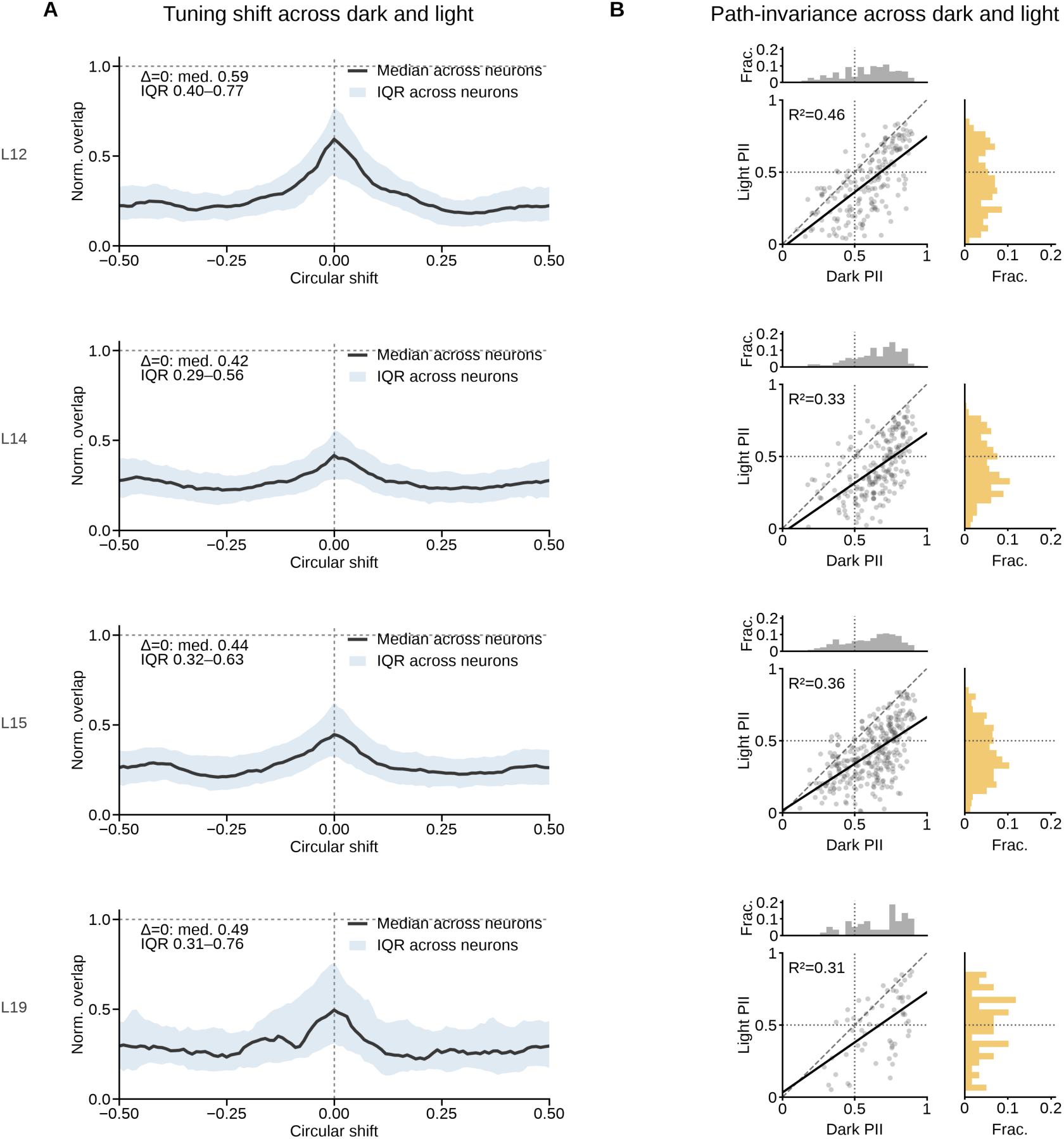
**(A)** Per-animal data of Fig. 2B. **(B)** Per-animal data of Fig. 2C.

**fig. S4.**
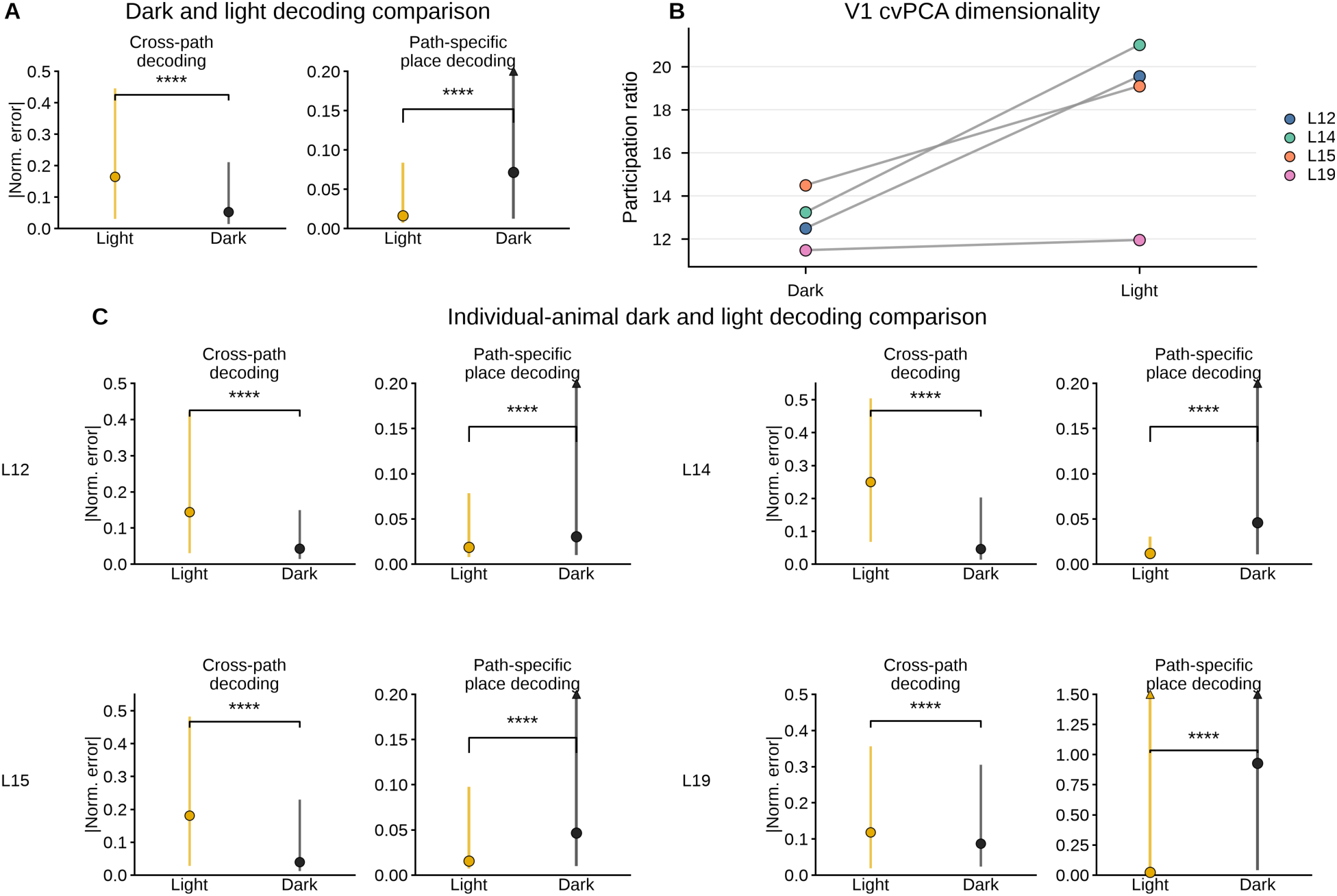
**(A)** Cross-path decoding of same-turn paths (left) and path-specific place decoding (right) in light and dark. Compared with light, cross-path decoding error is lower in darkness, whereas path-specific place decoding error is higher, consistent with light-induced reduction in path-invariance. **(B)** Paired comparison of V1 population dimensionality between dark and light epochs, quantified as the participation ratio of cross-validated PCA spectra computed from tuning curves across the four paths. Colors indicate different animals. **(C)** Per-animal data for panel A. In panel A, significance denotes the largest per-animal *p*-value when the effect direction was consistent across animals; panel C shows animal-specific tests. \**p*<0.05, \*\**p*<10^-2^, \*\*\**p*<10^-3^, \*\*\*\**p*<10^-4^.

**fig. S5.**
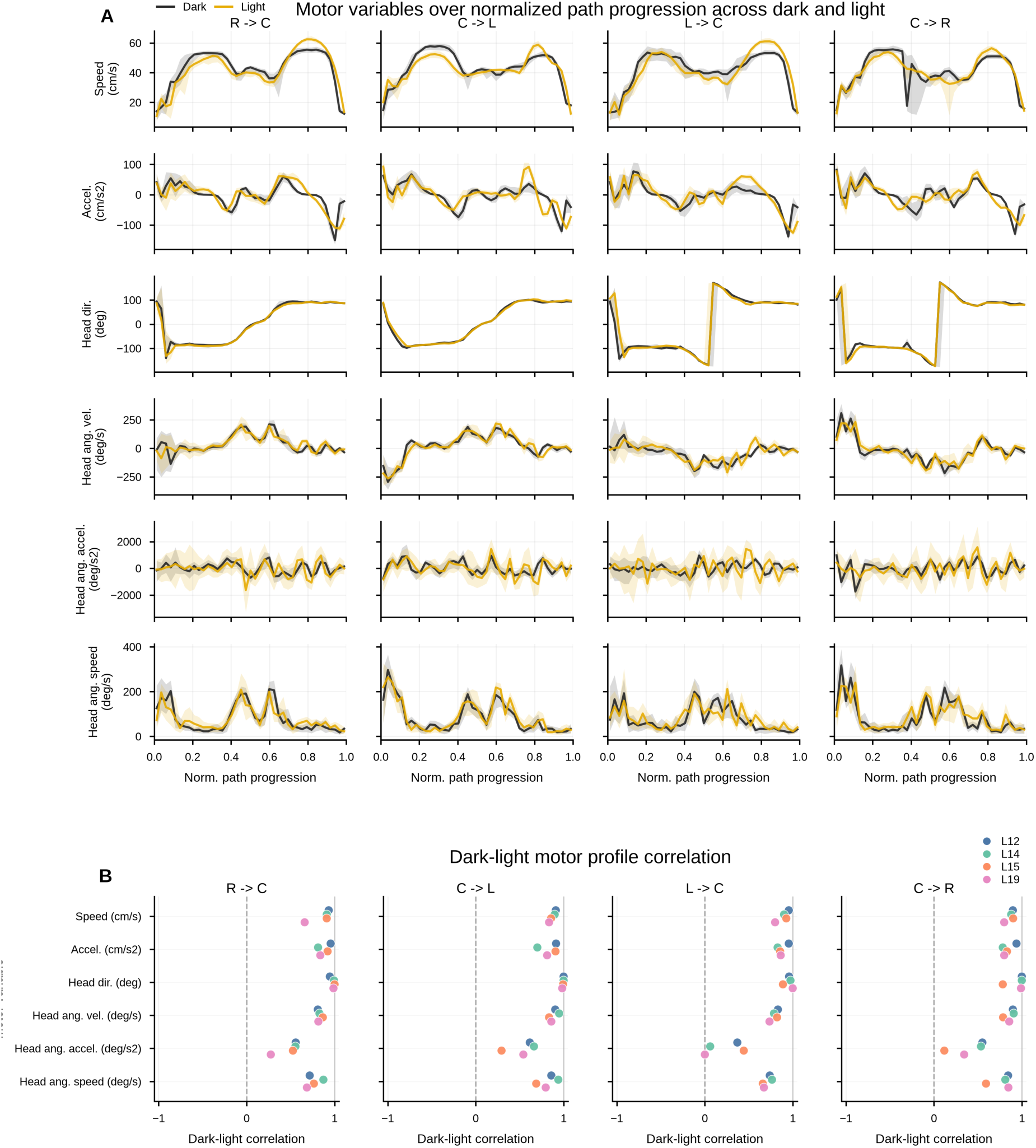
**(A)** Six motor variables aligned to normalized path progression for each of the four paths in light (yellow) and dark (black) epochs for a single animal (L14). Solid lines show medians and shaded bands show interquartile ranges across trials. **(B)** Correlation of the motor variables between light and dark epochs. Colors indicate different animals.

**fig. S6.**
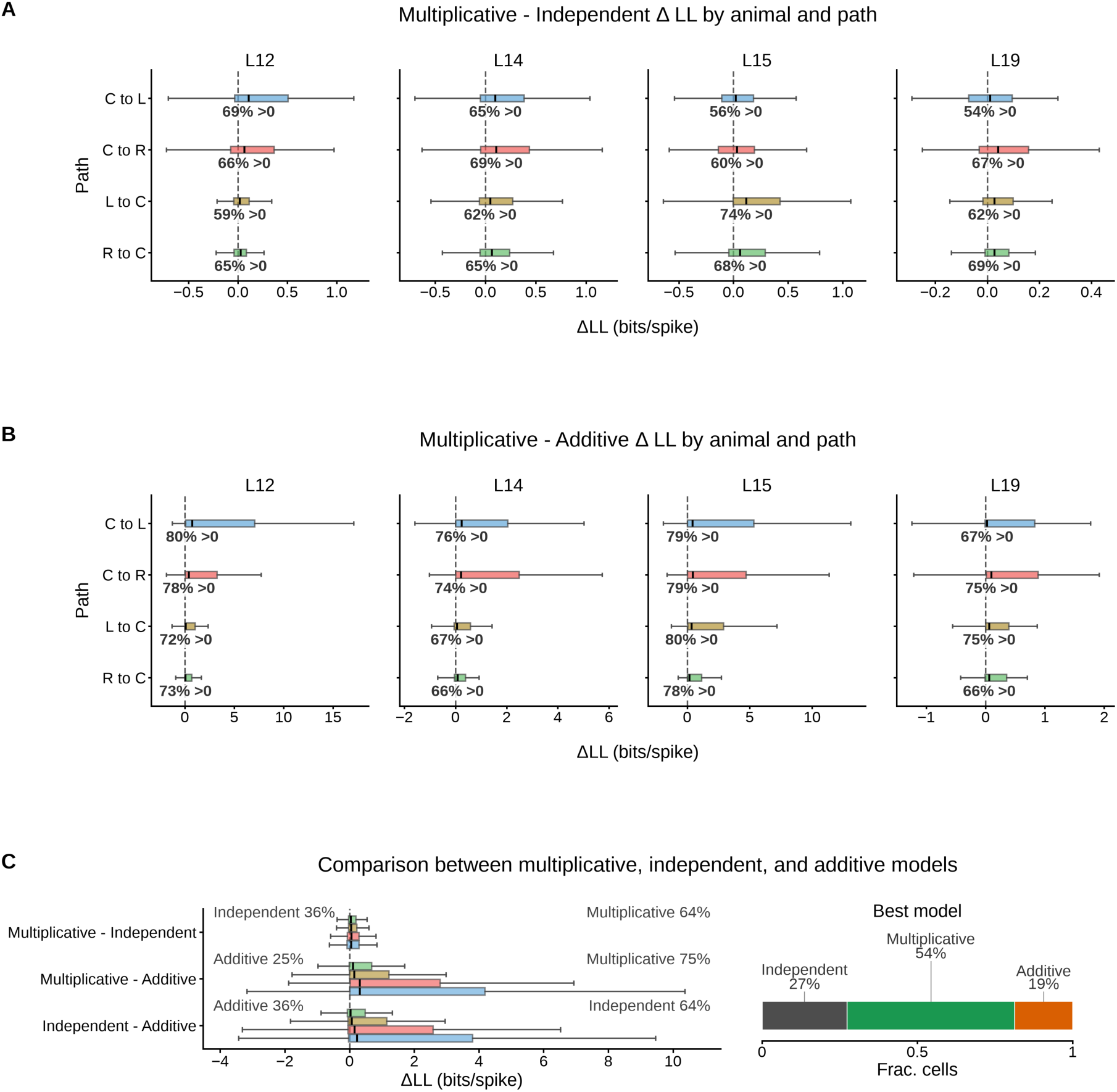
**(A)** Difference in log likelihood between the multiplicative and independent models (defined in Fig. 3A) for all four path types for each of the four animals used in this study (L12, L14, L15, and L19). Colors indicate different path types. The number to the right of each box-and-whisker plot indicates the fraction of cells with ΔLL greater than 0 (favoring multiplicative model). **(B)** Same as (A) but for multiplicative and additive models. **(C)** Left: Three-way comparison of pair-wise differences in log likelihood among multiplicative, independent, and additive models. Right: Fraction of V1 neurons best explained by each of the three models.

**fig. S7.**
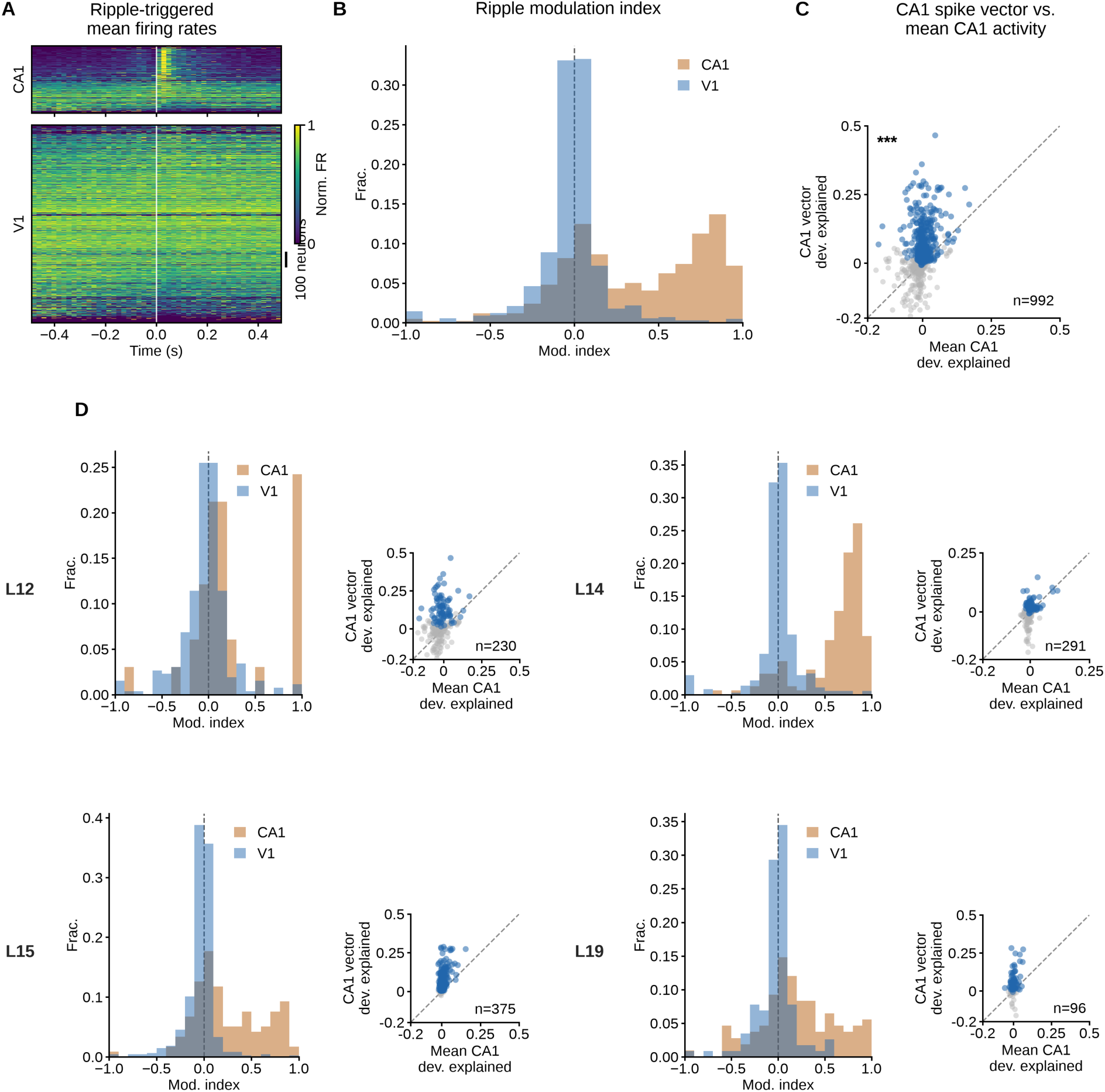
(**A**) Heatmap of mean ripple-triggered firing-rate across awake ripples. The modulation is very strong for CA1 neurons but much weaker for V1 neurons. **(B)** Distributions of ripple modulation indices for V1 (blue) and CA1 (orange) neurons during light epochs, pooled across sessions. For each neuron, the index was calculated as ((R-B)/(R+B)), where R and B are the mean peri-ripple firing rates from 0 to 100 ms and −500 to −300 ms relative to ripple onset, respectively. (**C**) Comparison of GLM prediction when using CA1 spike vectors vs. mean CA1 activity. Deviance explained for mean CA1 activity is mostly at 0 whereas CA1 vector-based prediction is much higher. The dashed line indicates equal performance; blue points denote units significantly predicted by the population-vector model in the shuffle test (*p*<0.05), and gray points denote nonsignificant units. Significance was assessed using a one-sided, 100,000- permutation linear mixed-effects test of the paired model difference with animal as a random effect. \*\*\**p*<10^-3^. **(D)** Per-animal data of panels B and C.

**fig. S8.**
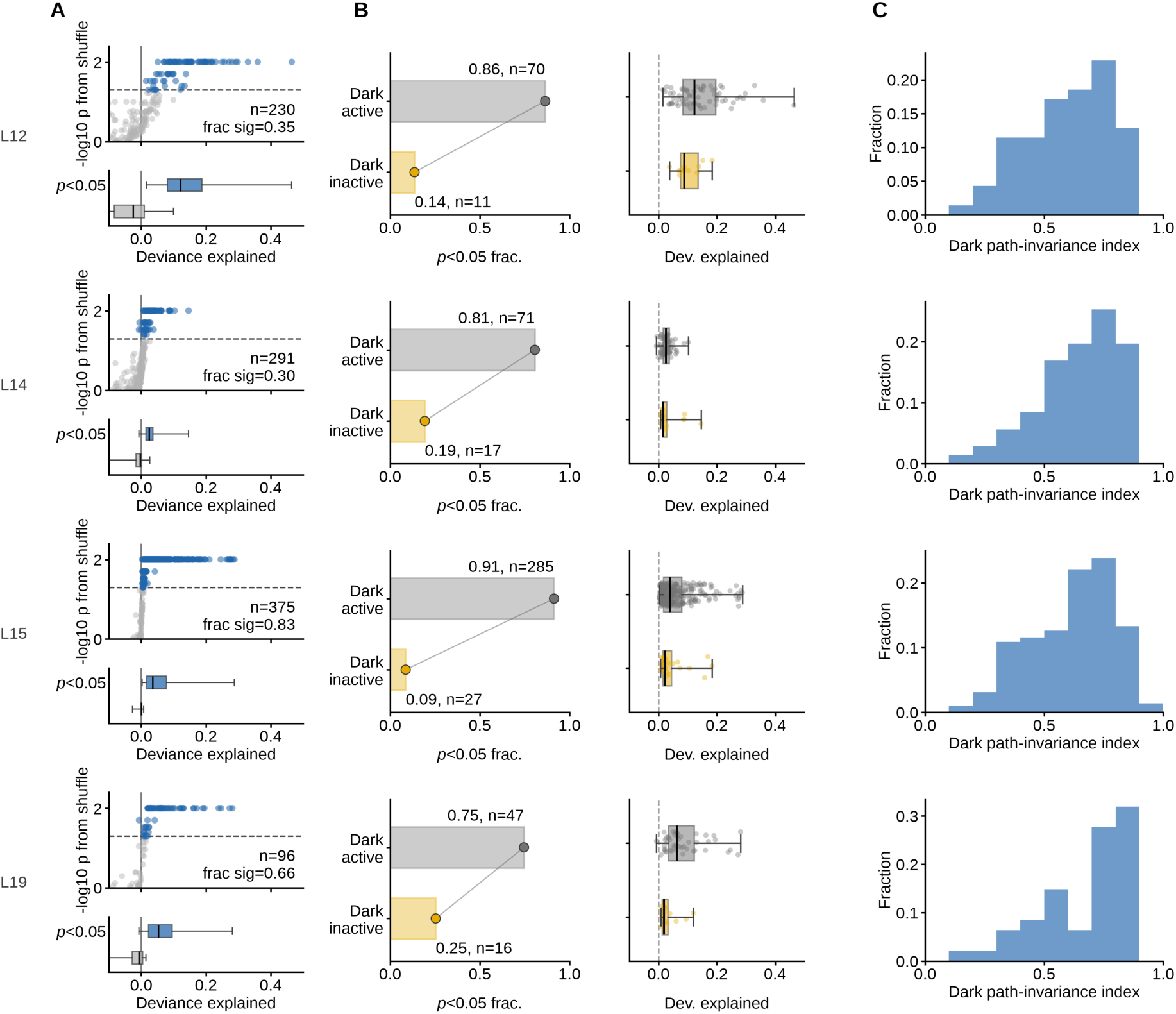
(**A**) Per-animal data of Fig. 4D. **(B)** Per-animal data of Fig. 4E. **(C)** Per-animal data of Fig. 4F.

## Materials and Methods

### Software

Electrophysiology and behavior data were collected with <u>Trodes</u> (version 2.4). Tracking of the animal’s body parts in the video recordings was done with <u>`DeepLabCut`</u> (version 2.3.7) (*39*). Data were converted to NWB format by <u>`pynwb`</u> (version 3.1.3) (*40*) and <u>`trodes_to_nwb`</u> (version 0.1.9). Spike sorting was done with <u>`spikeinterface`</u> (version 0.103.2) (*41*) and <u>`mountainsort4`</u> (version 1.0.7) (*42*). Construction of tuning curves and decoding analyses were done with <u>`pynapple`</u> (version 0.10.3) (*43*). Ripple detection was done with <u>`ripple`_`detection`</u> (version 1.7.1). GLM analyses were done with <u>`nemos`</u> (version 0.2.5). Analysis pipelines were constructed and the results were saved using <u>`Spyglass`</u> (version 0.5.5) (*44*). Visual stimuli were generated with <u>`psychopy`</u> (version 2023.1.2) (*45*). All analysis scripts are available on GitHub at https://github.com/khl02007/v1ca1. An AI coding assistant (GPT 5.5) was used to help with the implementation of the analysis scripts.

### Animals and surgical implantation

Four adult male Long–Evans rats were used in this study (4–10 months old; 400–600 g at implantation). All procedures were performed under protocols approved by the UCSF Institutional Animal Care and Use Committee.

Under anesthesia and analgesia and using aseptic techniques, bilateral craniotomies and durotomies were made over primary visual cortex (V1) and dorsal hippocampal area CA1. Each rat was implanted with four 128-channel flexible polyimide probes fabricated at Lawrence Livermore National Laboratory, with one probe targeting each hemisphere of V1 and CA1.

Target coordinates were AP −4.0 mm, ML ±2.0 mm, and DV 3.1 mm for CA1, and AP −8.0 mm, ML ±3.2 mm, and DV 1.8 mm for V1. AP and ML coordinates were referenced to bregma and the midline, respectively; DV denotes depth below the brain surface. Probes were typically implanted in the sagittal plane with the shanks parallel to the midline. Probe locations were verified histologically after the experiments (fig. S1A).

### Behavioral training and W-maze task

After food restriction to ∼85% of body weight, rats were first trained to alternate between the two ends of a linear track for sugary milk. Rats generally acquired the behavior (>80% trials correct) within one day, corresponding to approximately three 25-min epochs. They were then trained to perform spatial alternation on a W-maze (Fig. 1A). On outbound trials, rats were rewarded for alternating between the left and right outer wells; inbound trials consisted of returning from either outer well to the center well. This produced four correct path types: center-to-left, left-to-center, center-to-right, and right-to-center.

Each recording day comprised four to five 25-min run epochs separated by 25-min rest epochs. The visual condition was fixed within each run epoch and varied across epochs, including an initial stimulus configuration, a stimulus-swap configuration in which stimuli on the two outer arms were exchanged, and a dark condition. There was also a gray screen condition that was not analyzed for this manuscript. These visual manipulations were introduced from the first day of W-maze training. Rats typically reached greater than 90% correct performance after 2–3 days of training. We often collected data for another 7–10 days. The analyses reported here used data collected from well-trained animals (performance >90%) on training days 5–7.

### Visual environment and dark condition

Visual stimuli were presented on monitors (Asus PA278QV, Sceptre E205W) positioned 5–6 cm from the edges of the left and right outer arms and the horizontal segment of the W-maze. The center arm contained no monitors and was enclosed by walls covered with black corrugated plastic sheets. Gaps between adjacent monitors were covered with black plastic sheets to reduce uncontrolled visual cues. The stimuli included gray screen, sinusoidal gratings, black-and-white dots of various sizes (5–25° in visual angle) randomly positioned across the monitor, and black-and-white checkerboard. Visual stimuli remained fixed throughout each epoch. During stimulus- swap epochs, the stimulus configurations assigned to the left and right outer arms were exchanged.

During dark epochs, all monitors and visible-light sources were switched off or masked with black electrical tape, and behavior was recorded under near-infrared illumination with a peak wavelength of 940 nm. We measured the full emission spectrum (UDT Instruments, S471) and total irradiance (Thorlabs PM100 with S120C sensor) at maze level with the sensor oriented parallel to the track, which was ∼0.01 mW/cm². This irradiance was 48-fold below the 0.48 mW/cm², 930-nm illumination under which dark-adapted Long-Evans rats performed at chance in a high-contrast visual-form discrimination task, and approximately fourfold below the 0.0415 mW/cm², 850-nm illumination that rats failed to detect in an operant light-detection task (*28*, *29*). Thus, the illumination was below published irradiance levels from infrared sources that failed to support visually guided behavior in rats. Estimated rod activation was less than 10 R*/rod/s; this value represents a conservative upper bound rather than a behavioral visibility threshold.

### Position tracking and behavioral variables

Behavior was recorded using an overhead camera (Allied Vision, Manta G-158C). Video was acquired at 30 frames/s during light epochs and 15 frames/s during dark epochs, allowing a longer exposure time under infrared illumination. An infrared-reflective marker attached to the headstage was used to define head position and was tracked by DeepLabCut. The base of the neck and the base of the tail were also tracked by DeepLabCut. Head direction was calculated from the vector between the base of the neck and the headstage marker. Linear speed, acceleration, signed angular velocity, angular speed, and angular acceleration were calculated from the tracked 2D position after interpolation.

### Linearization of 2D position and definition of path progression (Figs. 1–3)

We linearized the 2D, DeepLabCut-tracked head position by projecting it onto a 1D graph that covers each path. This was done separately for each of the four path types for most analyses. Normalized path progression was then defined by dividing the linearized position by the length of the graph for one path, such that progression was 0 at the starting reward well and 1 at the ending reward well. For the allocentric-place model in Fig. 1F, the track linearization was done for the entire W-maze, in the following order: the center arm, the left horizontal segment and the left arm, and the right horizontal segment and the right arm.

### Tuning curve calculation and population heatmap (Fig. 1B–D, Fig. 2A)

During each dark epoch, position was linearized separately for the four directed W-maze paths and scaled to a normalized path-progression axis ranging from 0 to 1. Analyses of visual and path-related activity patterns were restricted to movement periods (speed >4 cm/s). Traversals of each path were divided into odd- and even-numbered trials, and occupancy-normalized firing-rate tuning curves were calculated using `pynapple` separately for the two subsets using 50 path-progression bins and Gaussian smoothing (σ = 1.5 bins). For each ordering path, neurons were sorted according to the peak location of their odd-trial tuning curves. This ordering was then applied to the held-out even-trial tuning curves for all four paths. Repeating this procedure with each path as the ordering reference produced the 4 × 4 heatmap matrix in Fig. 1D, in which rows indicate the path used for ordering and columns indicate the path displayed. For visualization, each neuron’s even-trial tuning curve was divided by its own peak firing rate separately for each path. V1 neurons were pooled across animals and included if their dark- epoch movement firing rate was at least 0.5 Hz and their odd–even tuning correlation was at least 0.5 for one or more paths. This latter criterion identified neurons that showed reliable firing in the task (800 of 1262 neurons met this criterion).

### Comparison with motor coding (Fig. 1E)

During movement in the dark epoch (speed >4 cm/s), spike counts were binned in 50-ms intervals. We fit two nested, ridge-regularized Poisson GLMs with a log link using `nemos`. The reduced model contained motor covariates only: running speed, linear acceleration, head angular velocity, head angular acceleration, absolute head angular velocity, and the sine and cosine of head direction. Motor covariates were derived from DeepLabCut head and body coordinates expressed in centimeters and evaluated in the same nonoverlapping 50-ms bins used for spike counts. Linear speed was calculated as the magnitude of the head-point velocity after Gaussian smoothing of the horizontal and vertical velocity components (σ = 0.1 s) and was linearly interpolated to the bin centers. Linear acceleration was calculated as the numerical time derivative of the binned speed. Head direction was defined as *θ* = atan2(*y*_head_ − *y*_body_, *x*_head_ −*x*_body_) and represented by sin(*θ*) and cos(*θ*) to account for its circularity. For angular derivatives, head direction was unwrapped before interpolation; signed angular velocity and angular acceleration were then obtained by successive numerical differentiation, and angular speed was defined as the absolute angular velocity. In total, six motor quantities (linear speed, linear acceleration, head direction, signed angular velocity, angular acceleration, and angular speed) produced seven model regressors (head direction contributed separate sine and cosine terms). For each cross-validation split, regressors were z-scored using the mean and standard deviation of the training data and the same transformation was applied to held-out data; no spline expansion of motor variables was used.

The full model contained the same motor covariates together with path-invariant progression terms comprising a turn-direction offset and separate fourth-order B-spline functions of path progression for left- and right-turn paths. Center-to-left and right-to-center paths shared the left-turn function, whereas center-to-right and left-to-center trajectories shared the right-turn function.

Models were evaluated using nested, trial-wise cross-validation. Trials from each of the four paths were divided separately among five outer folds. Within each outer training set, three-fold cross-validation was used to select the ridge penalty (10^-1^ to 10^-6^) independently for each model. For the full model, spatial resolution (2, 4, or 8 cm) that determined the number of spline basis functions was selected jointly with ridge penalty using cross-validated median information.

Held-out log likelihoods were summed across outer folds, and the contribution of path-invariant progression beyond motor variables was quantified as log likelihood difference in bits per spike:

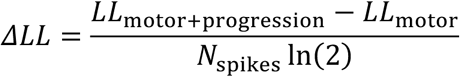

Positive values indicated that including path-invariant progression improved prediction beyond motor variables alone.

The analysis was restricted to V1 neurons with an odd–even spatial tuning curve correlation > 0.5 for at least one of the four path types. The >0.5-Hz movement-rate criterion was applied to the outer training laps in Fig. 1E.

### Comparison with alternative codes (Fig. 1F)

To distinguish path-invariant progression coding from allocentric place and distance-traveled coding, we compared three tuning-curve-based Poisson encoding models during movement in the dark epoch. Trials were assigned to five cross-validation folds separately for each path. For each training fold, we estimated occupancy-normalized firing-rate tuning curves in 4-cm bins and smoothed them with a Gaussian kernel with σ = 1 bin.

The path-invariant progression model used separate path-progression tuning curves for left- and right-turn paths, pooling the two paths associated with each turn direction. The allocentric-place model instead used a single direction-independent tuning curve defined on the linearized full W-maze. The distance-traveled model used a single normalized start-to-goal progression function shared across all four paths.

Held-out spike counts were evaluated in 50-ms bins using the firing rate predicted by the corresponding training-fold tuning curve. Log likelihoods were summed across folds, and each alternative model was compared with path-invariant progression as log likelihood difference in bits per spike:

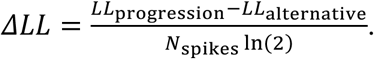

Positive values indicated better held-out prediction by path-invariant progression.

The analysis was restricted to V1 neurons with an odd–even spatial tuning curve correlation > 0.5 for at least one of the four path types. The >0.5-Hz movement-rate criterion was applied to the complete dark epoch in Fig. 1F.

### Decoding

We decoded one-dimensional spatial variables from V1 spiking using the Bayesian decoder implemented in `pynapple.decode_bayes`. The decoder used conditionally independent Poisson likelihoods and a uniform prior over spatial bins. The bin with the maximum posterior probability was taken as the decoded value. Spatial bins had a nominal width of 4 cm. Spikes were counted in 20-ms bins, and counts from four consecutive bins were summed using a rectangular sliding window, producing overlapping 80-ms decoding windows advanced every 20 ms. The true spatial coordinate was interpolated at each decoding timestamp, and decoding error was calculated as the absolute difference between the decoded and true coordinates. Tuning-curve estimation and decoding were restricted to movement periods.

#### Cross-path decoding (Fig. 1G; Fig. S2C; Fig. S4A,C)

Cross-path decoding tested whether V1 activity generalized between different path types. The decoded variable was normalized path progression, ranging from 0 to 1 along each trajectory. Thus, absolute error was already expressed as a fraction of one path’s length. For each directed transfer, path progression tuning curves were estimated from V1 spikes and position during traversals of one source path and then used to decode traversals of a different target path. V1 units with a mean movement firing rate greater than 0.5 Hz during the dark epoch were included. For the light–dark comparisons shown in Fig. S4A,C, the same population selected using dark-epoch firing rates was used in both epochs. Decoders were nevertheless fitted separately within the light and dark epochs. Fig. 1G and Fig. S2C included four directed transfers in each of three categories: same-turn/cross-arm, opposite-turn/same-arm, and matched inbound-or-outbound type/cross-arm. Fig. S4A,C included only the four same-turn/cross-arm transfers.

### Path-specific place decoding (Fig. S4A,C)

For path-specific place decoding, the four directed trajectories were assigned separate, nonoverlapping segments of a concatenated linearized coordinate, preserving both path identity and position along the path. Decoding was performed separately within each light or dark epoch using 5-fold trial-wise cross-validation. Trials were shuffled and assigned to folds separately for each trajectory; the training or held-out trials from all four trajectories were then pooled within each fold. All retained V1 units were included.

Absolute decoding error was divided by the length of one complete W-maze path.

#### Statistical test

For descriptive visualization, decoding errors were summarized using the median and interquartile range of all finite sample-level errors. Samples were pooled across animals for Fig. 1G and Fig. S4A and summarized separately for each animal in Fig. S2C and Fig. S4C.

Statistical inference used individual target-path traversals as the unit of analysis. For each traversal, we calculated the median absolute decoding error. Tests were conducted independently for each animal. The statistic was

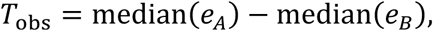

where *e_A_* and *e_B_* were the trial-level median errors for the two conditions. Condition labels were permuted within target-path strata while preserving the observed number of trials assigned to each condition. For each animal and comparison, 100,000 permutations were performed. Two-sided empirical *p*-values were calculated as

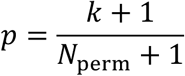

where *k* is the number of permutations in which |*T*_perm_| ≥ |*T*_obs_| and *N*_perm_ is the total number of permutations (100,000).

For Fig. 1G and Fig. S2C, same-turn/cross-arm errors were compared separately with opposite- turn/same-arm errors and matched-inbound-or-outbound/cross-arm errors. For Fig. S4A,C, light and dark errors were compared separately for cross-path and path-specific place decoding.

For the pooled significance annotations in Fig. 1G and Fig. S4A, the maximum two-sided *p*- value across the four animals was used. A significance symbol additionally required the expected effect direction in every animal: lower same-turn/cross-arm error in Fig. 1G, higher light than dark cross-path error in Fig. S4A, and lower light than dark path-specific place error in Fig. S4A. The annotations in the individual-animal panels, Fig. S2C and Fig. S4C, were instead derived from the corresponding animal’s own test.

### Shift in tuning across dark and light conditions (Fig. 2B)

For each V1 neuron, path progression tuning was computed separately for each of the four paths and for the light and dark epochs. Analyses were restricted to movement periods and tuning curves used bins corresponding to 4 cm of path length. A neuron–path pair was included only when its movement-period firing rate exceeded 0.5 Hz and its odd-versus-even-lap tuning correlation exceeded 0.5 for that same path in both conditions.

For each eligible neuron–path pair, missing tuning-curve bins were linearly interpolated, and the dark and light curves were independently normalized to unit area. This removed differences in overall firing-rate gain and isolated changes in tuning shape and position. Holding the dark curve fixed, we circularly shifted the light curve through every possible integer-bin displacement, with activity shifted beyond one end of the path wrapped to the other end. At each shift (Δ), overlap was calculated as

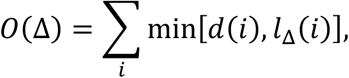

where *d*(*i*) and *l*_Δ_(*i*) are the unit-area dark and shifted-light tuning curves. Overlap ranged from 0 for nonoverlapping curves to 1 for identical curves. Shifts were expressed as signed fractions of the full path, from -0.5 to 0.5. Each exact periodic shift profile was then linearly interpolated, without smoothing, onto a common grid of 101 shifts.

To account for differences in tuning reliability among neurons and paths, overlap was rescaled separately for each neuron–path pair:

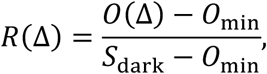

where *O*_min_ was the minimum overlap across all exact circular shifts and *S*_dark_ was the unit-area overlap between the odd- and even-trial dark tuning curves. Thus, *R* = 0 represents the least- overlapping circular alignment, whereas *R* = 1 represents the reproducibility of the neuron’s dark tuning across trial subsets. Values were not clipped and could therefore exceed 1. Pairs with an invalid split-half overlap or a nonpositive normalization denominator were excluded.

Valid path profiles were first averaged within each neuron so that neurons contributing more eligible paths did not receive greater weight. The plotted population curve is the median across neurons, and the shaded region is the interquartile range across neuron-level profiles. The annotation at zero shift reports the median and interquartile range. In total, the analysis included 2,212 neuron–path profiles from 766 V1 neurons across four animals.

### Path-invariance across dark and light conditions (Fig. 2C)

Path progression tuning curves were computed for each path in bins corresponding to 4 cm of path length. Analyses were restricted to movement periods during which speed exceeded 4 cm/s. For each neuron, overlap was calculated separately for the two pairs of paths involving the same turn direction: center-to-left with right-to-center for left turns and center-to-right with left-to-center for right turns. For two firing-rate curves *r*_1_(*b*) and *r*_2_(*b*), overlap was defined as:

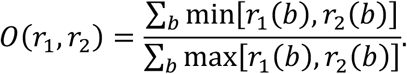

Thus, overlap retained differences in both tuning shape and absolute firing rate and ranged from 0, indicating no shared firing-rate area, to 1, indicating identical curves. Missing bins were linearly interpolated, and pairs with zero total firing-rate envelope were excluded. The path- invariance index was defined as the larger of the left- and right-turn overlaps. For paired dark– light comparisons, the turn direction with the larger overlap in darkness was selected for each neuron, and the overlap for that same direction was evaluated in the light epoch; the direction was not selected independently in light.

Path-invariance indices for V1 neurons across dark and light epochs were plotted in a scatterplot in Fig. 2C. Only V1 neurons with movement firing rates greater than 0.5 Hz in both epochs, finite overlaps in both conditions, and an odd–even tuning-curve correlation of at least 0.5 for at least one trajectory in each epoch were included. Neurons were pooled across animals, and the relationship between dark- and light-epoch values was summarized with an ordinary least-squares fit and its coefficient of determination (*R*^2^).

### Signal dimensionality from cross-validated PCA (fig. S4B)

To estimate the dimensionality of V1 path progression tuning, we analyzed one light (stimulus configuration AB) and one dark run from each of four sessions. Position was linearized separately for the four W-maze paths and divided into 4-cm bins. Only movement periods with speed greater than 4 cm/s were included. For each path and lighting condition, all trials were randomly assigned to four approximately equal, non-overlapping folds. Path progression tuning curves were calculated independently for each fold. Progression bins were retained only when occupancy was at least 0.01 s in every trial group under both lighting conditions, and the tuning curves from all four paths were concatenated.

Signal dimensionality was estimated separately in light and dark using four-fold cross-validated PCA. Tuning curves from three out of the four folds were averaged to form a training response matrix, while the remaining fold served as the held-out response matrix. The training and held-out matrices were independently mean-centered across progression bins. Each neuron was then scaled using its standard deviation in the training matrix. Principal axes were obtained from the training matrix by singular-value decomposition.

For each principal component *k*, reproducible signal variance was estimated as the covariance between the training and held-out projection scores:

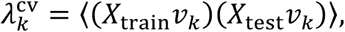

where *v_k_* is the training-derived principal axis and the average is across progression bins. Component-wise covariance estimates were averaged across the four held-out folds. Negative cross-validated spectral values were set to zero before dimensionality was calculated. Effective dimensionality was quantified using the participation ratio:

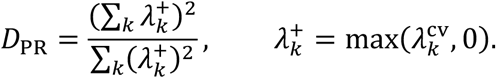

Higher participation ratios indicate that reproducible tuning variance was distributed across more population dimensions.

### Comparison of motor behavior in dark and light (fig. S5)

Head and body positions were used to calculate speed, acceleration, head direction, signed head angular velocity, head angular acceleration, and absolute head angular speed. Analyses were restricted to movement periods with speed >4 cm/s within traversals of four paths on the W-maze.

For fig. S5A, data from one animal (L14) were compared between one light epoch and one dark epoch. The motor variables were aligned to path progression bins. Lines show the median and shading shows the interquartile range. For fig. S5B, Pearson’s correlation was calculated from the median profile of the motor variables in light and dark epochs for the four animals included in this study.

### Sharp-wave ripple detection (Fig. 4)

Hippocampal sharp-wave ripples were detected from the CA1 recordings independently within each epoch from a prespecified set of 6–10 CA1 channels using the multichannel `Kay_ripple_detector` implemented in the Python package `ripple_detection` (v1.7.1). Wideband signals were band-pass filtered between 150 and 250 Hz using a fourth-order Butterworth filter applied in the forward and reverse directions and downsampled to 1 kHz. No notch filter was applied. For each channel, the analytic amplitude envelope was calculated, and a consensus ripple trace was constructed as the square root of the Gaussian-smoothed sum of squared envelopes across channels (σ = 4 ms). The consensus trace was z-scored using all valid samples within the corresponding epoch. Candidate ripples were defined as periods during which the consensus trace remained at least 2 SD above its mean for ≥15 ms; event boundaries were extended to the surrounding crossings of the mean. Closely spaced events were not additionally merged.

To exclude events during movement, speed was estimated from head position, smoothed with a Gaussian kernel (σ = 0.1 s), and interpolated to the ripple-LFP timestamps. Candidate events were retained only when speed at both event onset and offset was less than 4 cm/s.

### Models relating dark and light activity (Fig. 3, fig. S6)

For each V1 neuron and path type, spike counts in nonoverlapping time bins were modeled with ridge-regularized Poisson GLMs using a log link. Fits were restricted to movement periods with speed greater than 4 cm/s. Normalized path progression was represented using fourth-order B-spline basis functions, **B**(*p_t_*). A standardized linear speed term was included as a nuisance covariate and shared across lighting conditions.

#### Independent model

The independent model contained separate path-progression fields for the dark and light epochs:

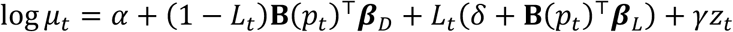

where *μ_t_* is the expected spike count, *p_t_* is normalized path progression, *L_t_* indicates the light condition, and *z_t_* is standardized speed. Thus, the light and dark spatial fields were estimated independently, without constraining their shapes to be related.

To generate a prediction on the held-out light epoch, the local light-field component within the exchanged segment was replaced by the corresponding component from the stimulus-matched source path.

#### Multiplicative model

The multiplicative model used a single path-progression field shared between dark and light epochs and modeled illumination-related changes as segment-specific gains:

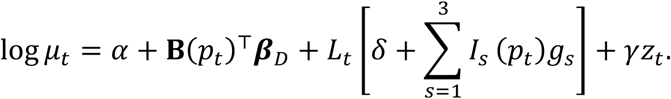

Here, *I_s_*(*p_t_*) denotes membership in one of three geometry-defined track segments. Because these terms entered through the log link, *g_s_* represented multiplicative firing-rate gains. The model therefore preserved the dark path progression tuning while allowing the firing rate to change independently within each path segment in light.

To generate a prediction on the held-out light epoch, the source path’s gain (including *δ*, which represents a shared component across the three segments) for that segment was transferred and combined with the target path’s dark tuning.

#### Additive model

The additive prediction was constructed from the fitted light and dark tuning curves from the independent model rather than fit as a separate GLM. For target path *q*, the prediction on the held-out light epoch was obtained by adding the light-minus-dark change measured on the stimulus-matched source path *q*′ to the target path’s dark tuning curve:

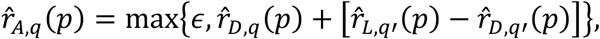

with *ε* = 10^−12^ spikes per bin (bin size: 50 ms) serving as the numerical floor.

#### Hyperparameters

Model hyperparameters were selected using 5-fold lap-level cross-validation. Candidate spline counts were 25, 40, and 60 and ridge strengths ranged from 10^-1^ to 10^-6^.

Candidate bin widths were 20 ms and 50 ms; 50 ms was selected in all cases. Spline complexity was selected using the median cross-validated information gain of the independent model across path–neuron combinations and was then shared across GLMs; ridge strength was selected separately for each model. The selected models were refit using all dark- and light-training movement data.

#### Quantifying prediction on stimulus-swapped held-out light epoch

The center-to-left path was paired with the center-to-right path at the terminal segment, and the left-to-center path was paired with the right-to-center path at the initial segment. All three models were scored on matched 50-ms spike counts from the exchanged segment using summed Poisson log likelihood; pairwise differences were reported in bits per spike.

### SWR-triggered mean firing rate and ripple modulation index (fig. S7A-B)

SWR-triggered firing rates were calculated for all CA1 and V1 units. For each unit, spikes occurring from 0.5 s before to 0.5 s after each ripple onset were counted in nonoverlapping 20- ms bins. Counts were averaged across SWRs within each recording epoch and divided by the bin duration to obtain firing rates. No temporal smoothing or baseline subtraction was applied. For visualization, each profile was divided by its maximum across the full 1-s window. Units were ordered separately within CA1 and V1 by decreasing SWR modulation index, defined as ((R- B)/(R+B)), where (R) was the mean firing rate 0–100 ms after SWR onset and (B) was the mean firing rate 500–300 ms before onset.

### CA1–V1 cross-correlograms (Fig. 4A)

CA1–V1 spike-time cross-correlograms were computed for the light epoch of one session (L15, light epoch with AB stimulus configuration). Spikes were restricted to all detected SWR intervals, and only units with at least 30 spikes during these intervals were retained. For every retained CA1–V1 pair, cross-correlograms were calculated using `compute_crosscorrelogram` in `pynapple` with CA1 as the reference population, 5-ms bins, and a nominal ±0.5-s lag window. The conditional V1 firing rate at each lag was divided by the V1 unit’s mean firing rate during the ripples; thus, values represent fold-change relative to ripple-period baseline, with 1 indicating baseline and positive lags indicating V1 spikes following CA1 spikes. Spikes were pooled across SWR intervals rather than calculating and then averaging a separate correlogram for each SWR.

For visualization, the three CA1 units with the strongest V1 partners were displayed. V1 units were grouped according to their relatively strongest displayed CA1 partner, ordered by peak three-bin mean cross-correlation, and the strongest one-third of each group was shown. Displayed curves were restricted to ±0.3 s, Gaussian-smoothed along the lag axis (σ = 5 ms), and clipped to the range 0–5. Each heatmap row represents one CA1-V1 pair.

### Predicting V1 from CA1 activity during ripples (Fig. 4B–D)

CA1-to-V1 coupling during SWRs was quantified separately for each run epoch using a ridge-regularized Poisson GLM. To avoid overlapping SWRs, only isolated SWR onsets separated from adjacent SWR onsets by at least 200 ms were retained. Each SWR constituted one sample: predictors were the vector of CA1 spike counts, and responses were the spike counts of individual V1 units, measured in the same 0–200-ms interval following ripple onset. V1 units were included if they emitted at least 0.1 spikes per SWR on average; all CA1 units were initially included as candidate predictors.

For V1 unit *j* and ripple *r*, the model was

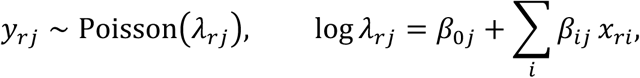

where *x_ri_* was the spike count of CA1 unit *i*. Models used a ridge-regularization strength of 0.1, fixed across sessions, and were optimized using the `JAXopts` L-BFGS solver in `nemos` (v. 0.2.5). Within each training fold, CA1 predictors with near-zero variance were removed; remaining predictors were standardized using training-fold statistics, divided by the square root of the number of retained predictors, and clipped to ±10. The same transformation was then applied to the held-out data.

Performance was evaluated using 5-fold cross-validation over chronologically ordered ripple samples, with each test fold forming a contiguous temporal block. For each V1 unit, Poisson deviance explained was calculated on held-out SWRs relative to a constant model whose expected count was estimated from the corresponding training fold. Deviance explained was then averaged across the five folds. The example observed-versus-predicted plots (Fig. 4C) show out-of-fold predictions.

Statistical significance was assessed using 100 response-shuffle refits per fold. For each shuffle, the training responses of each V1 unit were independently permuted across ripples, the GLM was refit, and performance was evaluated against the unchanged held-out responses. Observed and shuffled deviance-explained values were averaged across folds, and one-sided empirical *p*- values were calculated as

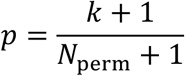

where *k* is the number of shuffles with performance on held-out responses at least as great as the actual performance and *N*_perm_ is the total number of shuffles (100). Units with *p*<0.05 were classified as significantly predicted.

### Predicting V1 activity with CA1 population vector vs. mean CA1 activity (fig. S**7**C)

To determine whether prediction of V1 activity depended on the activity of specific CA1 neurons rather than global CA1 activation, we fitted a reduced control model using the same SWRs, V1 units, and cross-validation folds as the full population-vector model. The reduced model replaced the vector of individual CA1 spike counts with the mean raw spike count across retained CA1 units during the 0–200-ms window following SWR onset. This scalar predictor was standardized using the training data separately within each fold. All other procedures were identical between models, including the Poisson GLM, ridge-regularization strength (0.1), 5-fold cross-validation, and calculation of held-out deviance explained relative to an intercept-only model estimated from the mean V1 response in the corresponding training fold.

Model performance was compared within V1 units using a one-sided permutation test based on a linear mixed-effects model. For each V1 unit with finite deviance-explained estimates from both models, we calculated the paired difference between the population-vector and mean-activity models. These differences were modeled as

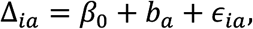

where *β*_O_ represents the population-level difference between models, *b_a_* is an animal-specific random intercept, and *ε_ia_* is the residual specific to animal *a* and unit *i*. The model was fitted by maximum likelihood. The one-sided alternative hypothesis was *β*_O_ > 0, indicating greater deviance explained by the population-vector model.

Statistical significance was evaluated using 100,000 paired permutations. For each permutation, the population-vector and mean-activity labels were independently exchanged within each V1 unit and the same mixed-effects model was refitted. The permutation *p*-value was calculated as

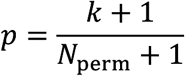

where *k* was the number of permuted coefficients greater than or equal to the observed coefficient and *N*_perm_ is the total number of permutations (100,000).

### Relationship between SWR predictability and dark-activity / path-invariance (Fig. 4E–F)

V1 units were classified as dark-active when their firing rate during movement in the dark epoch was greater than 0.5 Hz and as dark-inactive otherwise. We report the fraction of dark-active and dark-inactive cells among SWR-predictable units (*p*<0.05 in the vector-model shuffle test from Fig. 4D).

Enrichment of dark-active cells among SWR-predictable units was assessed using a linear-probability mixed-effects model, with dark-activity status as the binary response, SWR- predictability as a fixed effect, and animal as a random intercept:

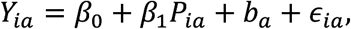

where *Y_ia_* is dark-activity status (0 or 1) and *P_ia_* is SWR-predictability (0 or 1) for animal *a* and unit *i*.

Significance was assessed by shuffling SWR-predictability labels within each animal while preserving the observed per-animal group sizes and refitting the model for each of 100,000 permutations.

Cross-validated deviance explained was compared between dark-active and dark-inactive SWR- predictable units using the linear mixed-effects model:

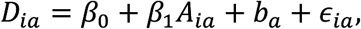

where *D_ia_* is cross-validated deviance explained and *A_ia_* is dark-activity status for animal *a* and unit *i*.

Dark-active labels were shuffled within each animal among SWR-predictable units while preserving group sizes, and the model was refitted for each of 100,000 permutations. Both models were fitted by maximum likelihood. Two-sided empirical *p*-values were calculated as

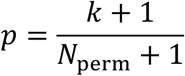

where *k* was the number of permutations with an absolute fixed-effect coefficient at least as large as the observed absolute coefficient, and *N*_perm_ is the total number of permutations.

Distributions are shown as box-and-whisker plots, with boxes indicating the median and interquartile range and whiskers spanning the full observed range.

Finally, we plotted the distribution of path-invariance index among SWR-predictable, dark-active V1 units. We compared the path-invariance index values in the predictable population against the non-predictable population using a linear mixed-effects model:

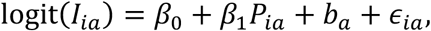

where *I_ia_* is the path-invariance index and *P_ia_* is SWR-predictability (0 or 1) for animal *a* and unit *i*. The model was fitted by maximum likelihood with animal as a random intercept. *I_ia_* was logit-transformed to avoid issues with fitting a bounded variable. Values were clipped to 10^−6^ and 1 − 10^−6^ prior to logit transformation.

Statistical significance was assessed by shuffling predictability labels within each animal while preserving the observed group sizes, refitting the model for each of 100,000 permutations, and comparing the absolute observed predictability coefficient with the permutation distribution. The two-sided empirical p-value was calculated as:

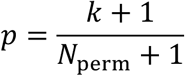

where *k* is the number of shuffled datasets with an effect at least as extreme as observed (u*β*^^^_1,perm_u ≥ u*β*^^^_1,obs_u) and *N*_perm_ is the total number of permutations (100,000).

## Acknowledgments

We thank Vanessa Bender for feedback on the manuscript and Viktor Kharazia for assisting with histology. We acknowledge GPT 5.5 for helping with editing parts of the text.

## Author contributions

Conceptualization: KHL, KK, MS, LMF

Methodology: KHL, JZ, JH, AY, RH, LMF

Investigation: KHL, PA, FG

Visualization: KHL

Funding acquisition: KHL, LMF

Project administration: KHL, MS, LMF

Supervision: KHL, KK, MS, LMF

Writing – original draft: KHL, LMF

Writing – review & editing: KHL, KK, MS, LMF

## Data, code, and materials availability

All code is available on GitHub (https://github.com/khl02007/v1ca1) and archived on Zenodo (https://doi.org/10.5281/zenodo.22155663). Raw and intermediate data are available as NWB files on DANDI (https://doi.org/10.48324/dandi.001958/0.260829.0404). A pre- populated Spyglass database is available as a Docker image (https://hub.docker.com/r/khl02007/spyglass-db-kyu_v1ca1) along with a corresponding Spyglass Hub image (https://hub.docker.com/r/khl02007/spyglass-hub-kyu_v1ca1).

